# The SV40 T-ag nuclear localization signal affects *Toxoplasma gondii* viability by targeting importin α

**DOI:** 10.1101/2025.07.20.665720

**Authors:** Manasi Bhambid, Janice Chithelen, Swati Patankar

## Abstract

Nucleocytoplasmic transport is essential in eukaryotes and mediated by importin (Imp) α/β receptors that recognize nuclear localization signals (NLSs) on cargo proteins. The SV40 large T-antigen NLS (SV40-NLS), studied extensively, binds importin α with high affinity in the nanomolar range, while modified versions, the Bimax peptides, bind in the picomolar range. Bimax peptides impede nuclear import and reduce viability in yeast and human cells, highlighting the potential of NLS peptides as inhibitors of nuclear transport. In this study, we investigated the potential of the SV40-NLS to target *Toxoplasma gondii* importin α (TgImpα). Expression of the SV40-NLS fused to a GFP reporter led to cytotoxicity in *T. gondii* tachyzoites; this depended on the SV40-NLS sequence and its position within the protein. Over-expression of TgImpα rescued parasites from the SV40-NLS-induced cytotoxicity, confirming that the mechanism of action involves disruption of nuclear import. Importantly, the same construct did not affect cell viability in mammalian cells, suggesting a selective vulnerability in importin α-mediated nuclear transport in the parasite. The SV40-NLS peptide offers an advantage over small-molecule inhibitors by targeting large interaction surfaces of importin α with high specificity, minimizing off-target effects. This study lays the groundwork for a novel peptide-based therapeutic strategy employing NLS motifs to selectively inhibit nuclear import in *T. gondii*.

## INTRODUCTION

Nucleocytoplasmic import in eukaryotic cells is mediated by nuclear transporters, importin α and β, that bind to nuclear localization signals (NLSs) on cargo proteins (Kalderon et al., 1984). The first identified NLS was found in the Simian Virus SV40 large tumour antigen (SV40 T-ag) (Lobl et al., 1990) and is extensively studied (Chang et al., 2012; Conti et al., 1998; Fanara et al., 2000; Harreman, Hodel, et al., 2003; Hodel et al., 2006; Hu & Jans, 1999). Subsequently, NLSs were identified in nuclear proteins and shown to bind the armadillo (ARM) repeats of importin α through a stretch of basic residues, mainly lysine and arginine; mutation of these basic residues decreases binding affinity to importin α and impacts nuclear targeting (Smith et al., 2017, 2018).

Interestingly, the N-terminal domain of importin α, the importin β-binding (IBB) domain, also contains NLS-like motifs and interacts with the NLS-binding sites in the ARM repeats (Kobe, 1999). These interactions result in auto-inhibition, preventing binding of the NLS on cargo proteins to importin α (Kobe, 1999). Auto-inhibition is relieved when the IBB domain interacts with importin β (Cingolani et al., 1999). This inhibitory function of the IBB domain regulates NLS-cargo binding and nuclear import (Harreman, Cohen, et al., 2003; Harreman, Hodel, et al., 2003). The strength of auto-inhibition varies: importin α proteins of *Homo sapiens*, *Mus musculus,* and *Saccharomyces cerevisiae* demonstrate strong auto-inhibition, while importin α proteins of apicomplexan parasites, *Toxoplasma gondii* and *Plasmodium falciparum,* show weak and lack auto-inhibition, respectively (Bhambid et al., 2023; Dey & Patankar, 2018; Harreman, Hodel, et al., 2003; Kobe, 1999; Miyatake et al., 2015).

Due to the essential roles of nuclear transporters, obstructing the import of nuclear proteins affects cell growth (Natarajan et al., 1996; Wang et al., 2010). Therefore, several small molecules targeting importin α in human cells have been identified; some of these molecules are in clinical trials for cancer and viral infections (Fraser et al., 2014; Kosyna & Depping, 2018; Tay et al., 2013; Thomas et al., 2018; Wagstaff et al., 2011, 2012). However, small molecules have multiple binding pockets and target proteins. In contrast, peptide inhibitors have the potential to bind to larger surfaces in target proteins, increasing specificity. Due to these features, peptide inhibitors have been widely proposed as potential therapeutic agents for many diseases (Banerjee et al., 2018; Fosgerau & Hoffmann, 2015; Lawrence et al., 2018; Wiedmann et al., 2017). These include inhibitors based on NLS peptides, showing a high specificity for importin α (Kosugi et al., 2008; W. Yu et al., 2018). For example, Bimax peptides, mutants of the SV40-NLS with picomolar binding affinities to *S. cerevisiae* importin α (ScImpα), cause loss of nuclear translocation and affect cell viability in yeast (Kosugi et al., 2008). Bimax peptides have been used to block nuclear transport in mammalian cells as well (Corona et al., 2013; Higa et al., 2018; Kosugi et al., 2008; Li et al., 2023; Marfori et al., 2012; Tunbak et al., 2016) and, due to their specificity for importin α, are commercially available (MedChemExpress).

Similarly, for the eukaryotic apicomplexans *T. gondii* and *P. falciparum,* which cause toxoplasmosis and malaria, and show increasing drug resistance and treatment failure (Montazeri et al., 2018; Siqueira-Neto et al., 2023), targeting importin α has been explored as a therapeutic strategy. Small molecules were identified through high-throughput screening that can inhibit importin α in biochemical assays, reduce nuclear transport and growth of the two parasites at multiple life cycle stages (Bhambid et al., 2025; Walunj, Dias, et al., 2022; Walunj, Wang, et al., 2022). While these studies demonstrate “proof-of-concept”, the small molecules exhibit IC_50_ values in the low micromolar range and affect host cell viability. This report explores the rationale of utilizing NLSs as highly specific peptide inhibitors of *T. gondii* importin α, resulting in the killing of the parasites.

We show that expressing SV40-NLS fused to a GFP reporter protein kills tachyzoites, a fast-replicating stage of *T. gondii*. Loss of viability was confirmed to be caused by the SV40-NLS, as healthy parasites were observed upon changing the sequence and position of the NLS. Parasites were rescued from the SV40-NLS toxicity when *T. gondii* importin α (TgImpα) was overexpressed, confirming the mechanism of action. Interestingly, numerous reports show the use of the SV40-NLS for carrying proteins into the nucleus in human cell lines, indicating that this NLS peptide does not have deleterious effects on the host cells of *T. gondii*. To confirm this, we performed SV40-NLS-GFP-His protein transfection in the Human Foreskin Fibroblast (HFF) and found no effect on viability. This report shows that, unlike results with Bimax peptides that bind strongly to yeast and human importin α, leading to loss of cell viability, wild-type SV40-NLS peptides hold potential in therapeutic targeting of *T. gondii*.

## MATERIALS AND METHODS

### Cloning of constructs

Primers used in the study are listed in Table 1 and were purchased from Sigma-Aldrich. Enzymes for cloning were obtained from New England Biolabs and Invitrogen (Thermo Fisher Scientific). All plasmids were confirmed by sequencing to be free of mutations.

**Table 1.**
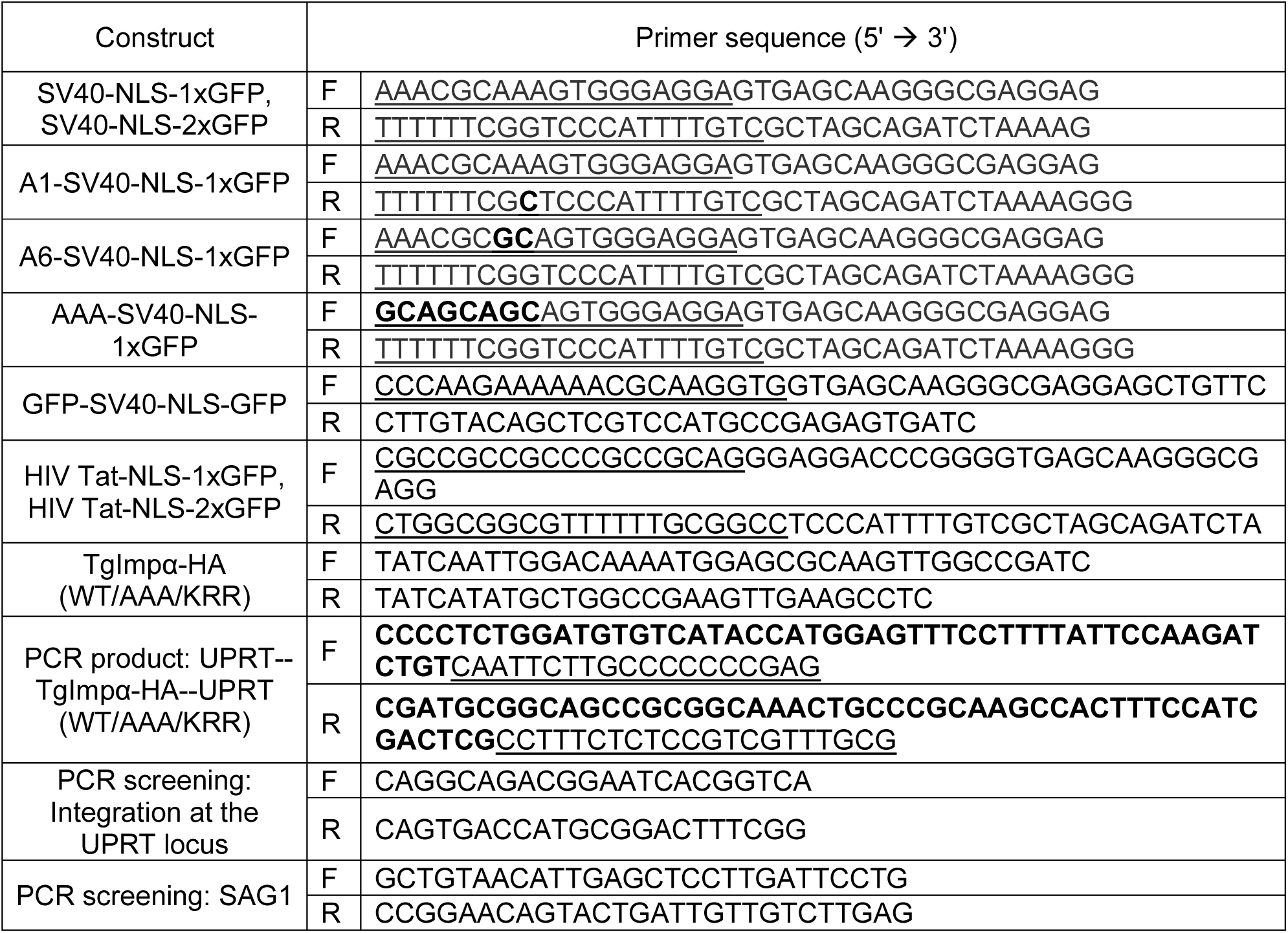
List of primers used in the study. The bases coding for the NLS motif (underlined) and the mutations (bold) are highlighted. For primers amplifying the pTubulin-TgImpα-HA-3’UTR PCR amplicon, the region homologous to the UPRT locus (bold) and the cassette to be inserted (underlined) are highlighted.

### NLS-GFP cloning

The wild-type SV40-NLS motif (PKKKRKV) was cloned N-terminal to GFP fragment in the pCTG-1xGFP and pCTG-2xGFP vectors (Bhambid et al., 2025). Expression was driven by the α-tubulin 1 promoter (ptub1) and regulated post-transcriptionally by the 3′ untranslated region (3′UTR) of the dihydrofolate reductase–thymidylate synthase gene (DHFR-TS). Alanine mutants, A1-SV40-NLS (**A**KKKRKV), A6-SV40-NLS (PKKKR**A**V), and AAA-SV40-NLS (PKK**AAA**V), were generated in the 1x-GFP vector. The NLS of the HIV type 1 transactivator of transcription protein (HIV Tat) is a cell-penetrating peptide (CPP) (Frankel & Pabo, 1988). The HIV Tat-NLS (GRKKRRQRRRPPQ) was cloned at the N-terminal of the GFP fragment in the pCTG-1xGFP and pCTG-2xGFP vectors.

### *T. gondii* importin α cloning

Wild-type *T. gondii* importin α (WT-TgImpα; TGME49_252290) was cloned in the pCTG-HA vector between the *MfeI* and *NdeI* restriction sites. The promoter and 3’UTR were the same as the pCTG-GFP vector. The TgImpα gene was PCR amplified from the WT-TgImpα-His plasmid (Bhambid et al., 2023). The mutants of the IBB domain, AAA mutant and KRR mutant constructs, were also cloned into the pCTG-HA vector with the same primers, generating AAA-TgImpα-HA and KRR-TgImpα-HA constructs.

### *T. gondii* importin α-HA integration into the genome

PCR products containing the TgImpα-HA expression cassette (wild-type and mutants) were integrated into the *T. gondii* genome at the uracil phosphoribosyl transferase (UPRT; TGME49_312480) locus using a CRISPR/Cas9 strategy. The CRISPR/Cas9 plasmid containing Cas9-SV40-NLS-GFP and a single-guide RNA for UPRT (UPRT-sgRNA) was utilized (Shen et al., 2014). Primers harbouring sequences homologous to 50 bp adjacent to the sgUPRT target were used to amplify the insertion cassettes.

### PCR screening

The integration of the TgImpα-HA cassette at the UPRT locus was confirmed by PCR. The forward primer was specific to the TgImpα cassette, while the reverse primer targeted the UPRT locus (Table 1). Successful genomic integration was indicated by the amplification of a 1483 bp product. The surface antigen 1 gene (*sag1*; TGME49_233460) was used as a housekeeping gene control (H. Yu et al., 2013).

### Culture maintenance and parasite transfection

*T. gondii* strains, RH and RHΔKu80, were maintained in Human Foreskin Fibroblast (HFF-1 SCRC-1041^TM^) cells (ATCC) and cultured in Dulbecco’s Modified Eagle Medium (DMEM; Invitrogen) supplemented with 10% Fetal Bovine Serum (FBS; HiMedia) and harvested as described (Striepen & Soldati, 2007). For transient transfection, parasites were harvested and electroporated as previously described (Bhambid et al., 2025). Either 50 μg of a single plasmid or 35 μg of each plasmid (for co-transfection of two plasmids) was used (Striepen & Soldati, 2007). The immunofluorescence assay was performed 22 hours post-transfection.

RHΔKu80 parasites were transfected with 50 μg of CRISPR/Cas9 plasmid and 10 μg of the ptub1-WT-TgImpα-HA-3’UTR amplicon. The AAA-TgImpα-HA and KRR-TgImpα-HA cassettes were transfected similarly. For stable transfection, the cuvette contents were subjected to two electroporation pulses. Transgenic parasite lines were cultured in DMEM supplemented with 1% FBS and selected using 9 μM 5-fluorodeoxyuridine (FUDR; MP Biomedicals) (Donald & Roos, 1995). Once parasite populations stabilized, clonal lineages were isolated with limiting dilution (Striepen & Soldati, 2007), and two clones from each line were selected.

### Protein transfection in HFF

Recombinant proteins GFP-His and SV40-NLS-GFP-His were purified as previously described (Dey & Patankar, 2018; Wagstaff & Jans, 2006; Walunj, Dias, et al., 2022). Protein concentration was determined using the Bradford assay. Proteins were incubated in 20 mM HEPES buffer at room temperature for 15 minutes (Chen et al., 2019) and then added to the HFF monolayer. The cells were incubated in DMEM (without FBS) for 24 hours at 37°C with 5% CO_2_.

### Immunofluorescence assay and microscopy

Immunofluorescence of the intracellular parasites expressing GFP-tagged constructs and HFF transfected with GFP-tagged proteins was performed as previously described (Bhambid et al., 2025). An additional antibody incubation step was included for parasites transfected with HA-tagged constructs. Samples were blocked with 3% Bovine Serum Albumin (BSA) in 1X PBS for one hour, then incubated with rabbit anti-HA primary antibody (1:500 dilution, CST) for two hours, followed by a goat anti-rabbit secondary antibody conjugated to Alexa 568 (1:400 dilution, Invitrogen) for one hour. Images were acquired by a Zeiss LSM 780 Confocal microscope with 100X objective or a Zeiss Observer Z1 Spinning Disc microscope with 40X or 63X objectives. To quantify GFP-positive parasitophorous vacuoles, 15 fields were imaged from each replicate in the Zeiss Observer Z1 Spinning Disc microscope at 40X.

### Image processing and analysis

The number of GFP-positive, HA-positive, and DAPI-positive parasitophorous vacuoles was counted manually, and the percentage of GFP vacuoles or HA vacuoles per DAPI counts in a field was calculated. A total of 45 fields were quantified per sample (15 fields per replicate performed in three independent experiments). Graphs were plotted using GraphPad Prism (version 8.4.3, San Diego), and the Mann-Whitney test was performed.

### Corrected total cell fluorescence (CTCF) analysis

Laser settings were kept constant during imaging for fluorescence intensity estimation and comparison. In ImageJ (version 2.0.0), a region of interest corresponding to the cell boundary was selected, and integrated density and area were quantified for both GFP and HA channels. An additional region of interest from the background was measured for background correction. The CTCF was calculated for each cell using the formula (El-Sharkawey, 2016).

CTCF = Integrated Density – (Area of selected cell X Mean fluorescence of background readings). Statistical analysis and data representation were performed with GraphPad Prism (version 8.4.3, San Diego).

### Western blots

The parasite pellet from a T-25 flask was added to lysis buffer (25 mM Tris-Cl pH 7.4, 150 mM NaCl, 1% TritonX-100, 1 mM EDTA, 5% glycerol, 2% protease inhibitor cocktail) and incubated on a rotatory shaker for an hour at 4°C. The Bradford assay determined protein concentration in the lysate. Protein loading dye was added to 80 μg of total protein, and the samples were heated at 100°C for 10 minutes before SDS-PAGE and overnight transfer to a PVDF membrane (Merck). The next day, the membrane was blocked with 3% BSA in 1X PBS-Tween 20 (PBST) for an hour, then incubated with rabbit anti-HA primary antibody (1:1000 dilution, CST) for 2 hours, followed by goat anti-rabbit secondary antibody conjugated to horseradish peroxidase (1:2000 dilution, GeNei) for an hour. The membrane was developed using chemiluminescent reagents and imaged using the Amersham ImageQuant 500 system. Subsequently, the membrane was washed thoroughly with 1X PBST, stained with Ponceau S, and imaged again. Band intensities were quantified using the Gel Analysis plugin in ImageJ (version 2.0.0) (Schneider et al., 2012). Protein expression was estimated by calculating the ratio of the TgImpα-HA band intensity in the chemiluminescence image to the total Ponceau-stained band intensity in the respective lane.

### Plaque assay and quantification

Confluent HFF monolayers in a 6-well plate were infected with 50 parasites/well. The plate was incubated undisturbed at 37°C, 5% CO_2_ for 7 days. On the seventh day, the monolayer was washed with 1X PBS, fixed with absolute ethanol for 5 minutes, and stained with Gram’s crystal violet solution (HiMedia) for 5 minutes. After washing and air drying, the plaques formed in the monolayer were imaged using a light microscope. For each plaque, the longest and the shortest diameter was measured using ImageJ (version 2.0.0). The plaque area was calculated using the formula (Foley & Remington, 1969; Jacot et al., 2020).

Plaque area = [(π X longest diameter X shortest diameter)/4].

Statistical analysis and data representation were performed with GraphPad Prism (version 8.4.3, San Diego).

### MTT assay

Proteins (GFP-His and SV40-NLS-GFP-His) were pre-incubated in 20 mM HEPES buffer as described earlier. Confluent HFF cells cultured in a 96-well plate were treated with increasing concentrations of these proteins for 24 hours. Cell viability was assessed using the MTT assay as previously described (Riss et al., 2004). Graphs were generated using GraphPad Prism (version 8.4.3, San Diego).

### Prediction of nuclear localization signal (NLS), nuclear export signal (NES), and post-translational modifications (PTMs)

The presence of a classical NLS (cNLS) and its score were determined by the cNLS mapper tool (https://nls-mapper.iab.keio.ac.jp/cgi-bin/NLS_Mapper_form.cgi) (Kosugi et al., 2008; Kosugi, Hasebe, Matsumura, et al., 2009). For determining the cNLS score of the NLS-GFP constructs used in this study, the input sequence was as follows: for example, MG**PKKKRKV**GG (NLS in bold) followed by the sequence of GFP as encoded in the pCTG-GFP vector.

Nuclear proteome datasets of *T. gondii* and *P. falciparum* (Barylyuk et al., 2020; Oehring et al., 2012) were taken and analyzed for the presence of cNLS by the cNLS mapper tool. For each protein, the highest-scoring monopartite cNLS sequence and its position were recorded and visualized using GraphPad Prism (version 8.4.3, San Diego). The proteins containing NLSs with a cNLS score ≥ 10.5 and positioned within the first 10 residues were further analyzed. Their predicted structure models were checked on the AlphaFold program (https://alphafold.ebi.ac.uk/) (Jumper et al., 2021; Varadi et al., 2022). The LocNES program (http://prodata.swmed.edu/LocNES/LocNES.php) (Simon-Areces et al., 2013) was used to predict the nuclear export signal (NES). Whether these proteins are post-translationally modified was checked on the phosphorylation, acetylation, and ubiquitination datasets available on ToxoDB (https://toxodb.org/toxo/app/) and PlasmoDB (https://plasmodb.org/plasmo/app) (Close et al., 2010; Cobbold et al., 2016; Green et al., 2020; Jeffers & Sullivan, 2012; Pease et al., 2013; Silmon de Monerri et al., 2015; Treeck et al., 2011).

## RESULTS

### Expression of the SV40-NLS in *T. gondii* tachyzoites results in a loss of viability

In a previous study (Bhambid et al., 2025), we reported that it was not possible to generate stable lines in *T. gondii* tachyzoites and *P. falciparum* blood stages using expression vectors where the SV40-NLS was fused at the N-terminus of GFP. To investigate this observation further, we transiently transfected *T. gondii* tachyzoites with plasmids expressing 1xGFP and SV40-NLS-1xGFP. Qualitatively, fewer vacuoles were observed expressing SV40-NLS-1xGFP compared to 1xGFP (Figure 1 a). Quantitative effects on parasite viability were analyzed by counting healthy, GFP-positive vacuoles in the population (Figure 1 b). While 24% of vacuoles were GFP-positive following 1xGFP transfection, only 12% were GFP-positive with SV40-NLS-1xGFP. This two-fold reduction indicates that expression of the SV40-NLS at the N-terminus negatively impacts parasite viability.

**Figure 1.**
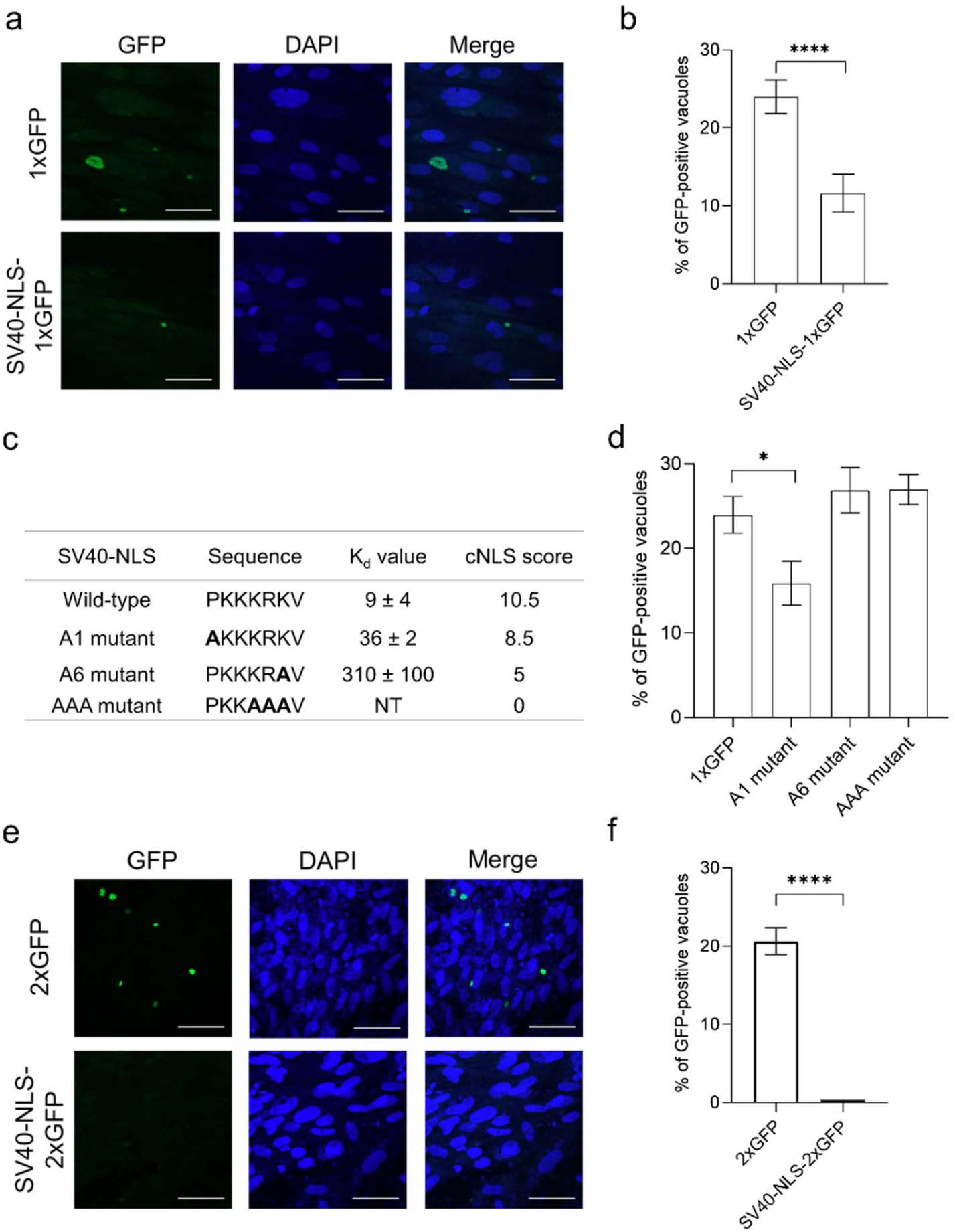
Loss of tachyzoite viability associated with SV40-NLS-2xGFP expression. **(a)** *T. gondii* tachyzoite vacuoles expressing 1xGFP construct alone and when fused with SV40-NLS. The images were captured on Zeiss observer Z1 Spinning Disc microscope with 40X objective (scale bar 50 μm); **(b)** Percentage of GFP-positive vacuoles per field were counted (N = 3) and the mean ± SEM was plotted; **(c)** The alanine mutations (bold) of the SV40-NLS motif generated in our study are shown, with their dissociation constant (K_d_) values (Hodel et al., 2006) and the cNLS score; **(d)** Percentage of GFP-positive vacuoles per field were counted (N = 3) with the mean ± SEM plotted; **(e)** Tachyzoite vacuoles expressing 2xGFP construct alone and when fused with SV40-NLS (scale bar 50 μm); **(f)** Percentage of GFP-positive vacuoles per field were counted (N = 3) and the mean ± SEM was plotted. A significant difference of p < 0.01 and p < 0.0001 is indicated by * and ****, respectively, on performing the Mann-Whitney test for all plotted graphs.

To confirm that the SV40-NLS sequence is responsible for the loss of parasite viability, mutants were generated. It is important to note that these mutants were selected based on a study that reported their affinity with ScImpα (Hodel et al., 2006); therefore, these mutants would be expected to exhibit lower affinities for TgImpα. Details of the three mutants, including amino acid sequences and binding affinities (K_d_ values), are given in Figure 1 c. The strengths of the wild-type SV40-NLS and the three mutants (A1, A6, and AAA) were predicted using the cNLS mapper tool (Kosugi, Hasebe, Matsumura, et al., 2009; Kosugi, Hasebe, Tomita, et al., 2009), which utilizes an algorithm based on the *S. cerevisiae* nuclear proteome. Based on these features, the strength of the wild-type NLS should be the highest (cNLS mapper score 10.5), with the strengths of the mutant NLSs being A1 > A6 > AAA (Figure 1 c).

Expression of the NLS mutants in tachyzoites showed a trend of increased GFP-positive vacuoles as the strength of NLS decreased (Figure 1 d), with mutants having weak NLS strengths (A6, AAA) showing comparable GFP-positive vacuoles to transfections expressing 1xGFP (∼25%). These results indicate that the effect of the NLS on parasite viability is linked to the SV40-NLS strength and can be correlated with binding to importin α, suggesting a mechanism of cytotoxicity.

Results shown thus far suggest that expression of the SV40-NLS hijacks a key protein of the nuclear transport machinery (TgImpα), resulting in a loss of parasite viability. We noticed that 1xGFP has a molecular weight below the published cut-off of the nuclear pore complex, allowing passive diffusion into the nucleus (Cardarelli et al., 2009; Timney et al., 2016). Reducing passive diffusion of SV40-NLS-1xGFP through nuclear pores by increasing the molecular weight of the reporter protein should result in increased cytoplasmic localization for interactions with TgImpα (Cardarelli et al., 2009). We hypothesized that this would, in turn, result in higher levels of cytotoxicity. The SV40-NLS was fused to a larger reporter, 2xGFP (Figure 1 e), and quantitative analysis showed 21% of vacuoles were GFP-positive upon 2xGFP transfection, while SV40-NLS-2xGFP expression resulted in complete cytotoxicity, *viz* 0% GFP-positive vacuoles (Figure 1 f).

Next, we assayed the cytotoxic effects of another NLS, a cell-permeable peptide (CPP) that can enter parasite and human cells, paving the way for future therapeutics. To this end, we selected the HIV Tat-NLS (GRKKRRQRRRPPQ) (Cardarelli et al., 2007; Smith et al., 2017), a CPP (Frankel & Pabo, 1988; Futaki, 2005; Morris et al., 2001; W. Yu et al., 2018) with a cNLS score of 8. The HIV Tat-NLS was fused to the N-terminus of 1xGFP and 2xGFP constructs, and upon transient transfection, we observed healthy, GFP-expressing vacuoles (Figure 2 a). Quantitative analysis revealed that approximately 17% and 19% of the population exhibited GFP-positive vacuoles in the HIV Tat-NLS-1xGFP and HIV Tat-NLS-2xGFP transfected populations, respectively (Figure 2 b). Despite the comparable cNLS scores of SV40-NLS and HIV Tat-NLS, and both are transported by importin α (Cardarelli et al., 2007; Smith et al., 2017), there are differences in the amino acid composition of the two NLSs (note the presence of arginines in the HIV Tat-NLS). These results highlight the uniqueness of the SV40-NLS in causing a loss of cell viability in tachyzoites.

**Figure 2.**
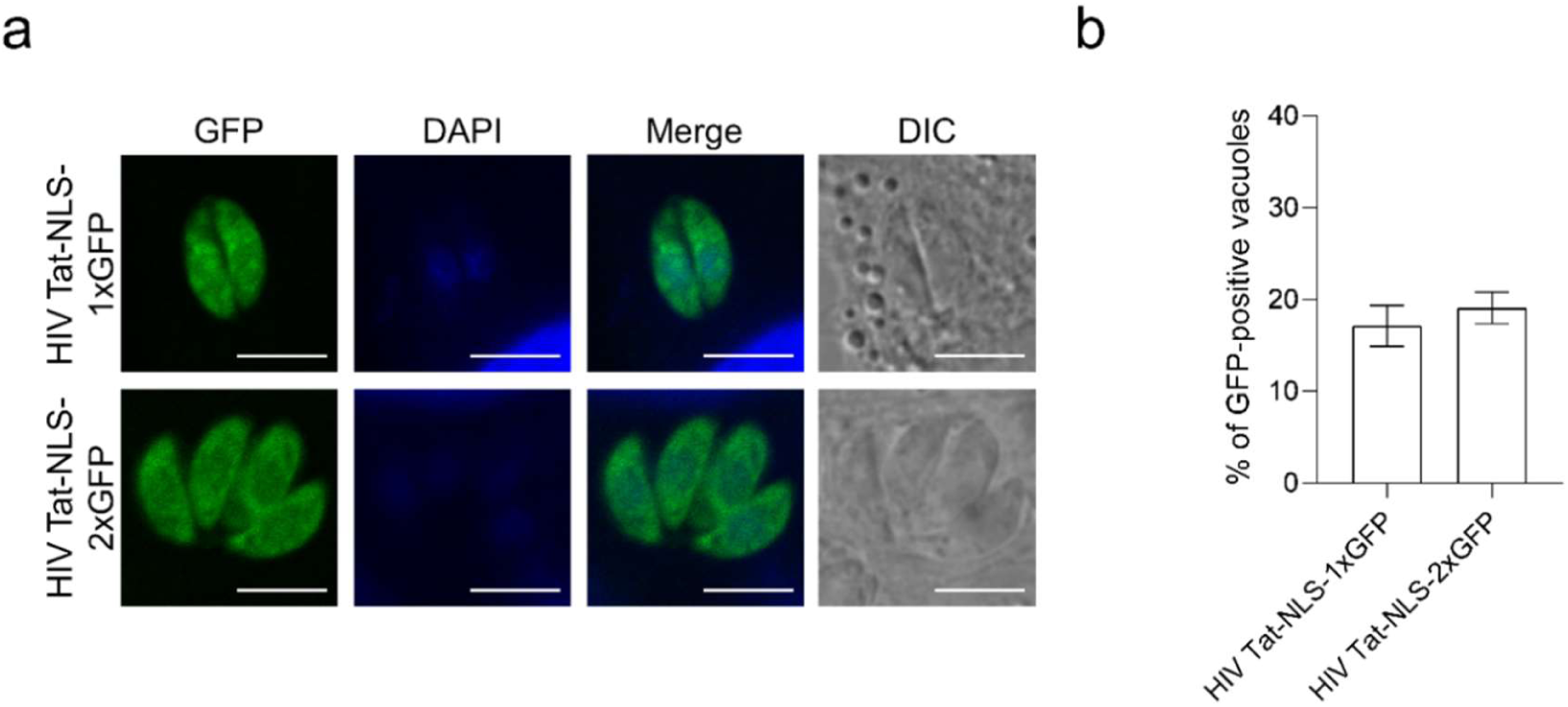
HIV Tat-NLS does not affect parasite fitness. **(a)** Localization of the HIV Tat-NLS-1xGFP and HIV Tat-NLS-2xGFP expressed in *T. gondii* tachyzoites. The images were captured on a Zeiss LSM 780 confocal microscope with a 100X objective (scale bar 5 μm); **(b)** Graph shows mean ± SEM of the percentage of vacuoles after transient co-transfection of GFP-positive vacuoles (N = 3). The Mann-Whitney test performed shows no significant difference.

Together, these results demonstrate that the SV40-NLS is the primary determinant of parasite cytotoxicity, with its severity dependent on both its binding affinity for TgImpα and the ability of the fusion protein to engage the nuclear transport pathway. This mechanism could explain our unpublished results showing an inability to generate stable lines expressing SV40-NLS-GFP in both *T. gondii* and *P. falciparum*.

#### Analysis of the nuclear proteome of apicomplexan parasites *P. falciparum* and *T. gondii*

As the SV40-NLS with a cNLS score 10.5 affects parasite viability, we wondered whether NLSs as strong as the SV40-NLS exist in the *T. gondii* and *P. falciparum* nuclear proteomes. An analysis of the strength of NLSs in apicomplexan nuclear proteins (identified by proteomics) has not been reported in the literature. Therefore, nuclear proteome datasets of *T. gondii* ME49 and *P. falciparum* 3D7 strains (Barylyuk et al., 2020; Oehring et al., 2012) were analyzed using the cNLS mapper tool, and the predicted monopartite NLS sequences, with associated score and location, were noted (Figure 3 a, Supplementary Data). We chose to assess the strength of the NLS by its cNLS mapper score, as this correlated well with the phenotype of loss of viability of parasites (Figure 1 c, d).

**Figure 3.**
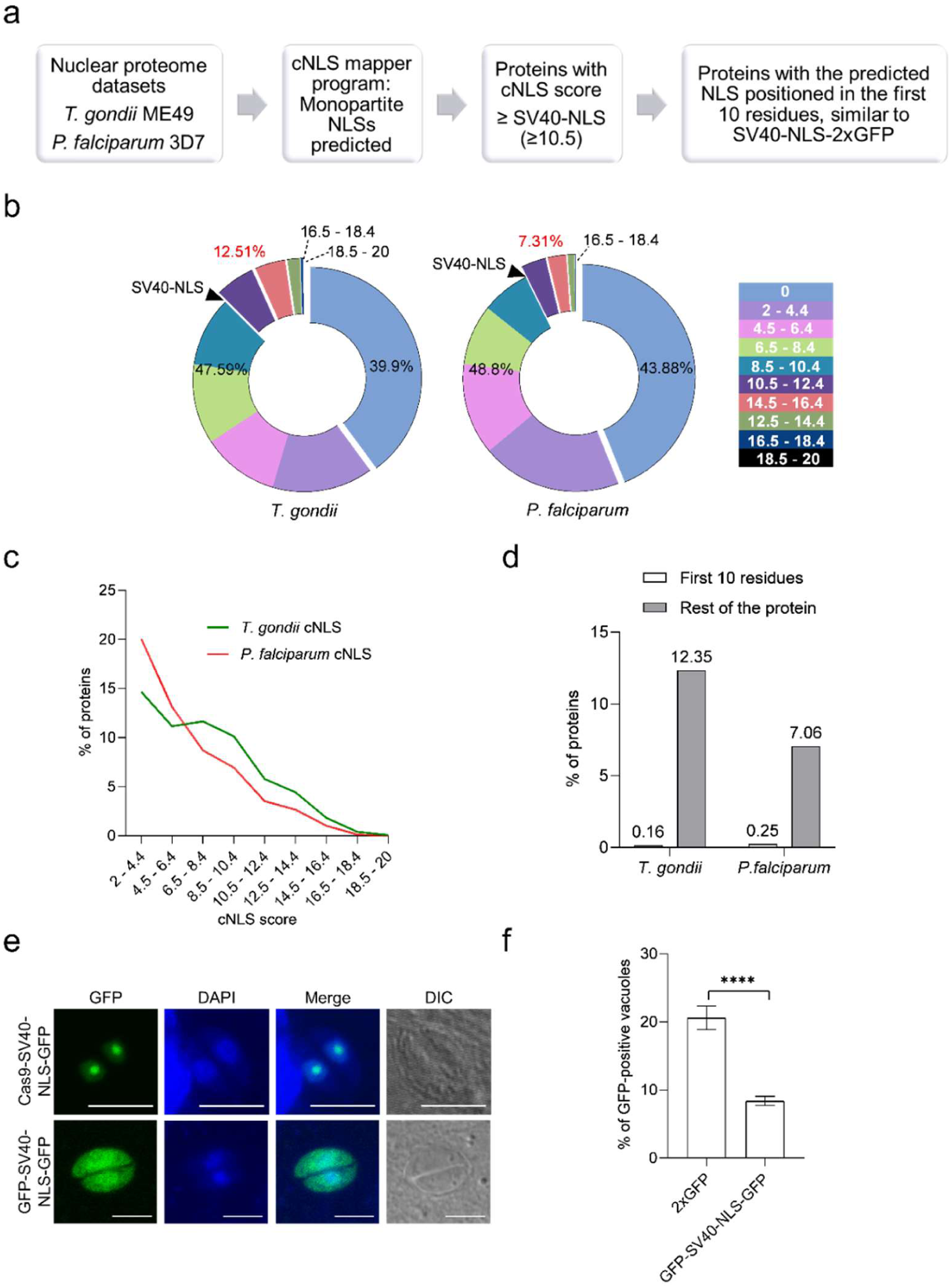
NLSs equal to and stronger than SV40-NLS exist in the *Toxoplasma* and *Plasmodium* nuclear proteomes. **(a)** Flow chart for the bioinformatic analysis performed on the nuclear proteome datasets of *T. gondii* and *P. falciparum*; Using the cNLS program, the predicted cNLS were classified according to their score in a **(b)** pie chart and **(c)** line plot. The arrow indicates where the SV40-NLS cNLS score (10.5) lies; **(d)** Graph shows the position of the predicted cNLS motif within the protein for those with a cNLS score ≥ 10.5; **(e)** Localization of Cas9-SV40-NLS-GFP and GFP-SV40-NLS-GFP in *T. gondii* tachyzoites. Images were captured in a Zeiss LSM 780 Confocal microscope with 100X objective (scale bar 5 μm); **(f)** Percentage of GFP-positive vacuoles per field was counted (N = 3), and the mean ± SEM was plotted. Mann-Whitney test shows a significant difference of p < 0.0001, indicated by ****.

In about 40% of the nuclear proteome, no cNLSs were predicted, suggesting that these proteins may not be strictly nuclear, may harbour non-classical NLSs, or be trafficked by other nuclear transporters (Harel & Forbes, 2004). Similar results in *P. falciparum* have been reported earlier (Oehring et al., 2012). Nevertheless, the program predicted cNLSs in more than half of the nuclear proteome, with scores ranging from 2 to 19 in both datasets (Figure 3 b). On distributing these scores into bins, we observed an inverse trend between the percentage of proteins and the cNLS score, with subtle differences between the two parasite datasets (Figure 3 c). Relatively, *T. gondii* harbours more proteins with strong cNLSs, and *P. falciparum* harbours more proteins with weaker cNLSs. It is tempting to speculate that the strength of cNLSs can be correlated with the strength of auto-inhibition, which is weak in the case of TgImpα and lacking in the case of PfImpα (Bhambid et al., 2023; Dey & Patankar, 2018).

Noteably, 12.51% and 7.31% of proteins in *T. gondii* and *P. falciparum* harbour NLSs stronger than the SV40-NLS (Figure 3 a, b). We further checked the location of these strong NLS motifs, particularly whether they are located in the first 10 amino acids (Figure 3 a), similar to our expression constructs, which fused the SV40-NLS at the N-terminal of GFP. Less than 0.3% of nuclear proteins were predicted to contain NLSs with a score ≥ 10.5, located within the first 10 amino acids (Figure 3 d). The proteins that fall into this category are ATPase, AAA family protein (TGME49_247390), cell-cycle-associated protein kinase PRP4 (TGME49_313180), cleavage and polyadenylation specificity factor subunit 4 (PF3D7_1419900), and 40S ribosomal protein S3 (PF3D7_1465900) (Supplementary Data). In order to understand why these endogenous NLSs with high cNLS scores may not impact viability similar to SV40-NLS, we checked parameters that could influence importin binding and nuclear import: (1) structure of the NLS motif in AlphaFold, (2) presence and structure of a nuclear export signal (NES), and (3) post-translational modifications in the NLS. Both *Plasmodium* proteins contained post-translational modifications in the predicted NLS, which might affect binding to importin α and a predicted NES. For the *Toxoplasma* proteins, we speculate that other reasons, such as low protein expression, might result in an inability to hijack the cellular importin α pool.

A previous study demonstrated that the position of the SV40-NLS motif (N- or C-terminal) affects its ability to interact with importin α (Wagstaff & Jans, 2006). As the bioinformatics analysis showed that NLS strength and position vary considerably in the nuclear proteome, we asked whether changing the position of the SV40-NLS in the 2xGFP reporter construct would affect cell viability. We cloned the NLS between the two GFP genes (Bhambid et al., 2025). We also tested a fusion protein extensively used in the genetic manipulation of *Toxoplasma* (Cas9-SV40-NLS-GFP) (Markus et al., 2019; Shen et al., 2014). Consistent with our hypothesis, healthy GFP-positive parasites were observed upon transfection of parasites with both constructs (Figure 3 e). While the GFP-SV40-NLS-GFP protein showed nuclear and cytoplasmic localization, the Cas9-SV40-NLS-GFP protein showed what appeared to be nucleolar localization. Compared to ∼20% GFP-positive parasites observed with the 2xGFP transfections, 10% GFP-positive parasites were observed when the NLS was fused between the two GFP reporters (Figure 3 f). While these data indicate that the GFP-SV40-NLS-GFP does show cytotoxicity, it is less than the 100% cytotoxicity observed when the SV40-NLS was fused at the N-terminus of 2xGFP (Figure 1 f). Clearly, the position of the NLS affects its ability to kill parasites, possibly due to changes in accessibility for importin α binding.

### SV40-NLS cytotoxicity occurs *via* nuclear import

SV40-NLS mutants, with reduced binding affinity to importin α, do not affect *T. gondii* viability; therefore, the cytotoxicity associated with the wild-type NLS is likely due to its interaction with *T. gondii* importin α. This binding affinity of the wild-type SV40-NLS to TgImpα, which exhibits weak auto-inhibition, may disrupt nuclear import dynamics, resulting in impaired parasite viability. Notably, our unpublished data, where we could not generate stable lines of the SV40-NLS-GFP in *P. falciparum,* is consistent with this hypothesis, as PfImpα lacks auto-inhibition.

If this hypothesis is correct, expressing TgImpα from a second copy should rescue the parasites from NLS-mediated cytotoxicity. For these experiments, we used HA-tagged TgImpα variants with differing levels of auto-inhibition: WT-TgImpα (weak auto-inhibition), AAA-TgImpα (lacking auto-inhibition), and KRR-TgImpα (stronger auto-inhibition) (Bhambid et al., 2023). All HA-tagged importin α proteins localized to nuclear and cytoplasmic compartments, with an intense signal observed in the nucleus (Figure 4 a). When co-transfection of the three TgImpα-HA variants with SV40-NLS-2xGFP was performed in transient transfections, viable GFP-positive vacuoles were observed 22 hours post-transfection. These GFP proteins colocalized with HA fluorescence across all importin α variants (Figure 4 b), where an additional copy of TgImpα, regardless of the strength of auto-inhibition, rescued the toxicity mediated by the SV40-NLS.

**Figure 4.**
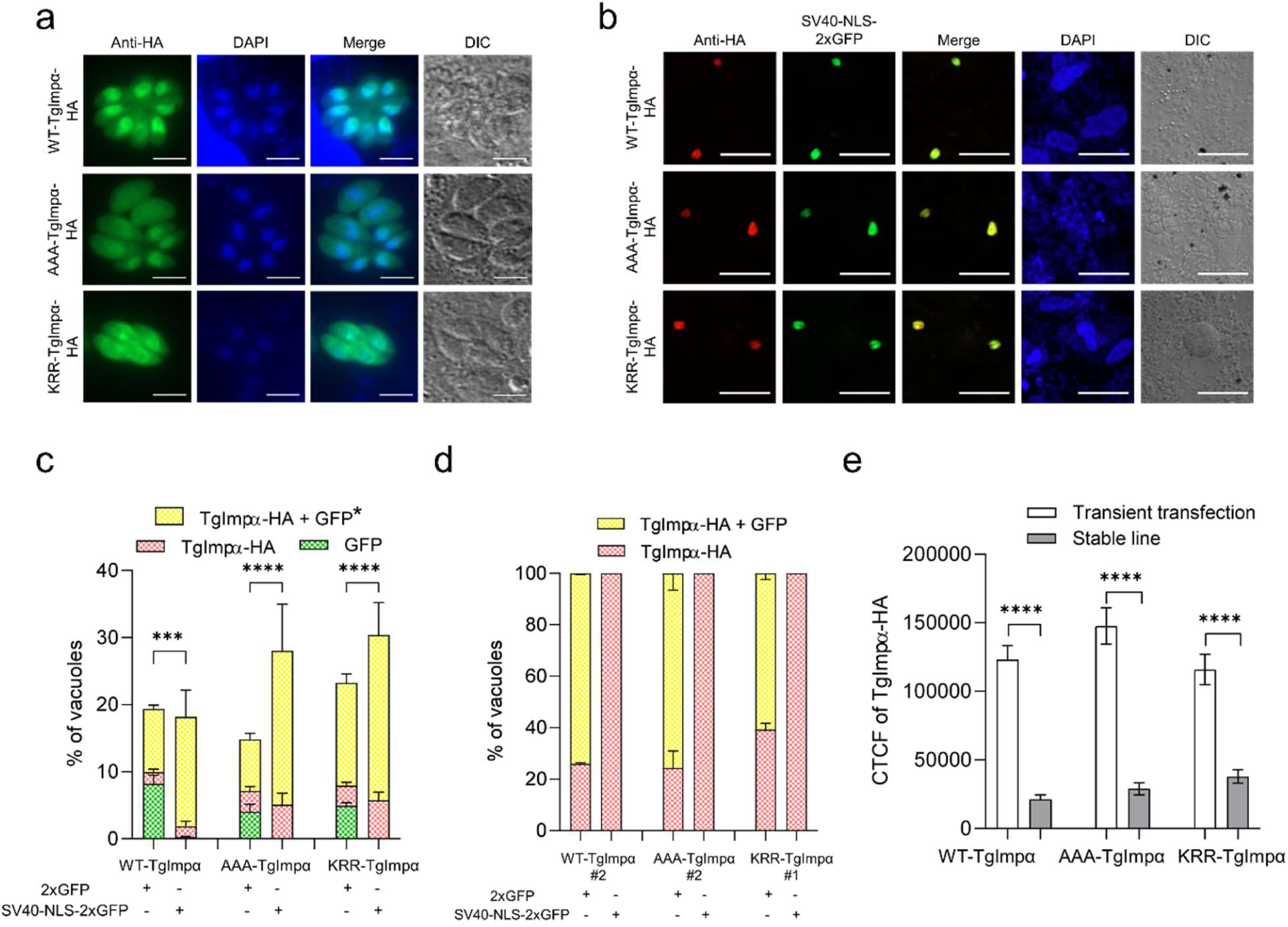
Overexpressing TgImpα protein rescues parasites from SV40-NLS-2xGFP toxicity. **(a)** WT-TgImpα-HA, AAA-TgImpα-HA, and KRR-TgImpα-HA localize in the nucleus and cytoplasm of *T. gondii* tachyzoites. Images were captured in a Zeiss LSM 780 Confocal microscope with 100X objective (scale bar 5 μm); **(b)** Transient co-transfection of WT and variants of TgImpα-HA (red) with SV40-NLS-2xGFP (green) resulted in live SV40-NLS-GFP expressing vacuoles. Images were captured in Zeiss Observer Z1 Spinning Disc microscope with 40X objective (scale bar 30 μm); **(c)** Graph shows mean ± SEM of the percentage of vacuoles after transient co-transfection expressing only SV40-NLS-2xGFP or 2xGFP (green), only TgImpα-HA (red) or both (yellow) per DAPI counts (N = 3). Mann-Whitney test performed shows a significant difference of p < 0.003 (***) or p < 0.0001 (****) for the yellow bar in the plotted graph; **(d)** Graph shows mean ± SEM of the percentage of vacuoles in the stable lines of TgImpα-HA (wild-type and variants) transfected with SV40-NLS-2xGFP or 2xGFP (N = 2); **(e)** Graph shows mean ± SEM of the CTCF intensity estimated of TgImpα-HA in transiently transfected and stable lines for 30-40 vacuoles. The Mann-Whitney test performed shows a significant difference of p < 0.0001.

The percentage of vacuoles expressing only SV40-NLS-2xGFP or 2xGFP (green), only TgImpα-HA, wild type or variants (red), or both (yellow) was quantified. When 2xGFP was co-transfected with wild-type or variants of TgImpα-HA, a green signal was observed in a population of vacuoles (Figure 4 c). When SV40-NLS-2xGFP was co-transfected with wild-type or variants of TgImpα-HA, 0% vacuoles expressed only green fluorescence, confirming that expression of this construct alone affects parasite viability (Figure 4 c). We next quantified the proportion of vacuoles co-expressing GFP and TgImpα-HA (yellow) and compared this proportion between the 2xGFP and SV40-NLS-2xGFP transfections. The proportion of vacuoles with yellow fluorescence showed a statistically significant increase in the case of SV40-NLS-2xGFP compared to those with 2xGFP, in the three TgImpα-HA transfections (Figure 4 c). These results show that overexpression of TgImpα-HA indeed rescues parasites from the cytotoxicity caused by the SV40-NLS, irrespective of the strength of auto-inhibition of TgImpα-HA. This finding confirms that the SV40-NLS-induced cytotoxicity is via the importin α-dependent nuclear transport pathway and that increasing the cellular pool of TgImpα can mitigate the detrimental effects of the NLS.

To further study the rescue of SV40-NLS-induced toxicity, we generated stable parasite lines constitutively expressing TgImpα-HA wild type and the variants showing different strengths of auto-inhibition. The ptub1-TgImpα-HA-3’UTR cassette was integrated into the UPRT locus using a CRISPR/Cas9-based strategy, followed by selection with FUDR (Shen et al., 2014) (Supplementary Figure 1 a). Single-plaque clones were screened for correct gene integration by PCR and protein expression by western blot (Supplementary Figure 1 b, c). Protein expression levels were quantified using Ponceau S-stained loading controls, and clones with the highest expression for TgImpα-HA and the two variants were selected for further analysis. No significant difference in growth rate was observed between the parental strain and the stable lines, as determined by plaque assay (Supplementary Figure 1 d). These results show that an extra copy of TgImpα does not affect the growth of tachyzoites in standard culture conditions. Immunofluorescence confirmed that HA-tagged proteins were distributed throughout the parasites in these clonal populations (Supplementary Figure 2 a).

These TgImpα-HA stable lines were transiently transfected with 2xGFP or SV40-NLS-2xGFP constructs (Supplementary Figure 2 b). When 2xGFP was expressed, a significant proportion of vacuoles were positive for both GFP and TgImpα-HA (yellow); no green vacuoles were seen, since all parasites express TgImpα-HA (Figure 4 d). However, when the SV40-NLS-2xGFP plasmid was transfected, no vacuoles positive for both GFP and TgImpα-HA were observed (Figure 4 d). Instead, the entire population was positive only for TgImpα-HA (red), showing a stark difference from the data from transient transfections (Figure 4 c). As no SV40-NLS-GFP-expressing vacuoles were observed, the stable lines did not show rescue of NLS cytotoxicity. It is important to note that in the transient transfections, immunofluorescence was performed at the same time point (22 hours), and here, TgImpα-HA was able to rescue NLS cytotoxicity. Quantification of the corrected total cellular fluorescence (CTCF) of the HA signal in transiently and stably transfected vacuoles revealed lower fluorescence intensity by orders of magnitude in the stable lines despite being driven by the same promoter (Figure 4 e). These results suggest that the concentration of importin α is a key factor in rescuing parasites from NLS-mediated cytotoxicity. This rescue appears to be independent of the auto-inhibition state of importin α.

### The SV40-NLS does not affect the viability of human host cells

The nuclear transport machinery is highly conserved between the human host and apicomplexan parasites; therefore, any NLS-based peptide therapeutic must be selectively tailored to target *Toxoplasma* and *Plasmodium* parasites without adversely affecting the host. Previous studies have examined SV40-NLS nuclear localization in various cells (Cardarelli et al., 2009; Hodel et al., 2006; Lyssenko et al., 2007). In one report, the localization of SV40-NLS-1xGFP and SV40-NLS-2xGFP constructs in HeLa cells was studied without assessing their impact on cell viability (Cardarelli et al., 2009). Another study reports that these constructs did not impair viability in yeast and human cells, but detailed viability data for human cells were not provided (Kosugi et al., 2008).

In light of these findings and to address this gap, we assessed the viability of human foreskin fibroblast (HFF) cells, used as hosts in our study, on transfection with GFP-His and SV40-NLS-GFP-His recombinant proteins (Dey & Patankar, 2018; Wagstaff & Jans, 2006; Walunj, Dias, et al., 2022). As the SV40-NLS motif lacks intrinsic cell-penetrating capability, we performed the HEPES protein-transfection method for intracellular delivery (Chen et al., 2019). GFP and SV40-NLS-GFP protein signals were detected within HFF cells at 7 μg/ml concentration, confirming successful protein uptake (Figure 5 a). The observed punctate localization patterns of both proteins suggest internalization via endocytosis and vesicle formation, consistent with previous reports on NLS-peptides and NLS-proteins (Chen et al., 2019; Cunningham et al., 2003; Koo et al., 2014; W. Yu et al., 2018). Additionally, with increasing concentration of proteins, a higher fluorescence signal was noted within the cell, as indicated by the CTCF values (Figure 5 b). While the host cells showed robust uptake of proteins, tachyzoites were not fluorescently labeled using this protocol (data not shown), indicating that alternate strategies must be used to deliver peptides into parasites. Nevertheless, cell viability of the host cells, assessed by MTT assay, indicated no cytotoxic effects of GFP-His and SV40-NLS-GFP-His proteins on the HFF cells across the tested concentration range (Figure 5 c). These findings indicate that SV40-NLS-GFP is well-tolerated by host cells while exhibiting cytotoxicity toward *T. gondii* tachyzoites.

**Figure 5.**
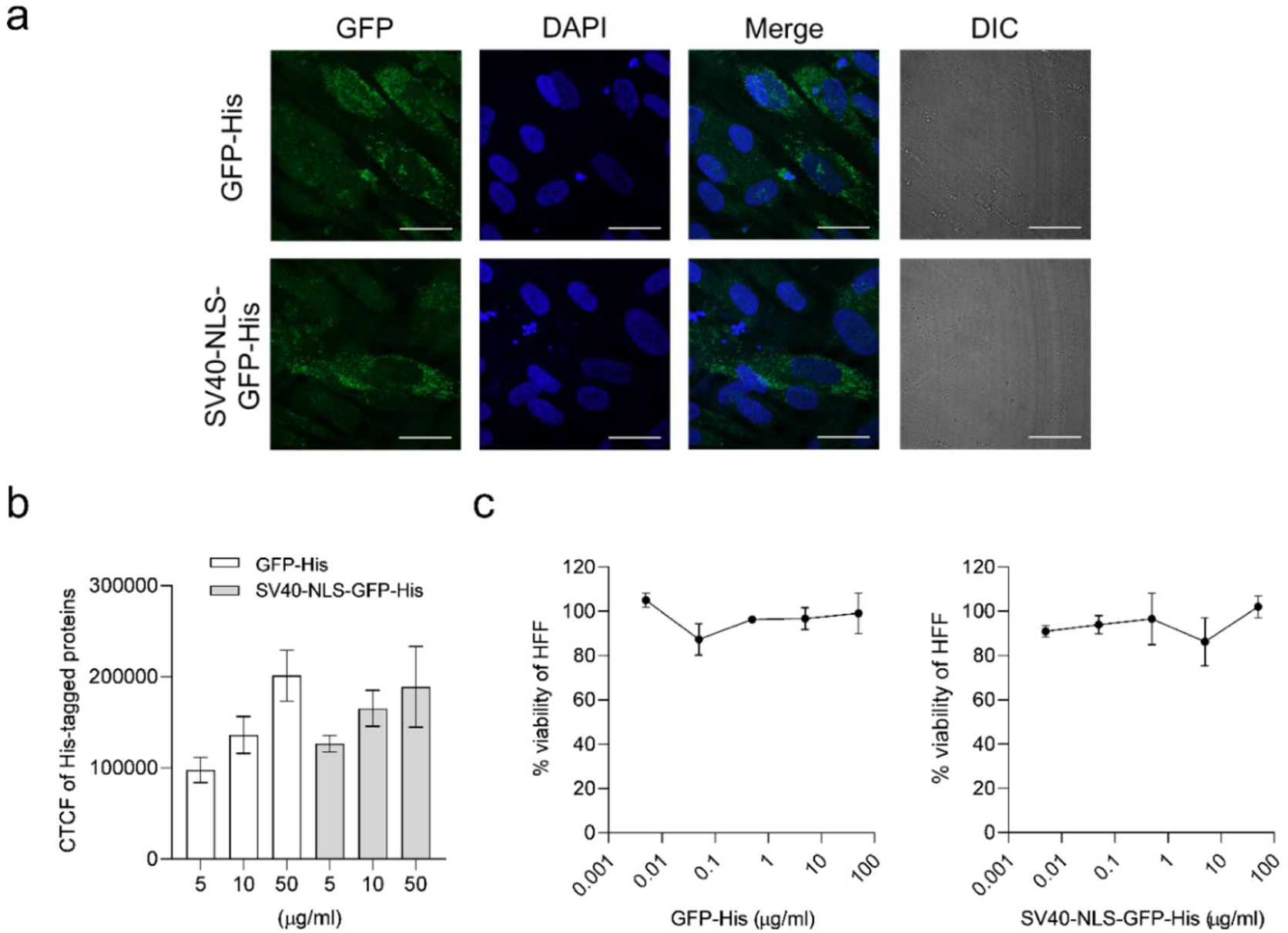
No effect of SV40-NLS-GFP on host cell viability. **(a)** GFP-His and SV40-NLS-GFP-His recombinant proteins (14 μg/ml) were transfected into HFF cells and incubated for 24 hours. The images were captured on Zeiss observer Z1 Spinning Disc microscope with 63X objective (scale bar 30 μm); **(b)** Graph shows mean ± SEM of the CTCF intensity estimated of GFP in 30 fields at varying concentrations of proteins (N = 2); **(c)** Cell viability MTT assay performed on HFF when treated with the proteins in the respective concentration range (N = 3).

## DISCUSSION

The emergence of drug resistance highlights the urgent need for new inhibitory molecules to combat diseases caused by apicomplexan parasites. Compounds with novel mechanisms of action are particularly valuable, as they may be effective against drug-sensitive and drug-resistant strains. One promising druggable target is the importin α protein, where small molecules inhibit binding to NLSs (Walunj, Dias, et al., 2022; Walunj, Wang, et al., 2022), parasite growth and stage differentiation in *P. falciparum* and *T. gondii* (Bhambid et al., 2025). This study proposes a novel inhibitory approach to target importin α in *T. gondii*.

### The SV40-NLS kills *T. gondii* tachyzoites via the nuclear import pathway

Expression of the SV40-NLS-2xGFP construct in *T. gondii* tachyzoites resulted in a pronounced cytotoxic effect. This appears to be a property of the sequence of the SV40-NLS and its location at the exposed N-terminus of the 2xGFP fusion protein. Indeed, an analysis of all predicted NLSs from the nuclear proteome of *P. falciparum* and *T. gondii* showed the presence of NLSs with scores much higher than that of the SV40-NLS. However, only a handful were found at the N-terminus of the endogenous proteins. While the expression levels of these proteins are unknown, apicomplexan parasites can tolerate strong NLSs in their proteomes.

We attribute the cytotoxic effect of the SV40-NLS to its unique sequence, resulting in a strong interaction with the TgImpα protein, potentially leading to a collapse of the nuclear import pathway. Evidence supporting this conclusion is based on the following findings: (1) altering the sequence or location of the SV40-NLS in the protein reduces toxicity, (2) another viral NLS (HIV Tat) does not cause cytotoxicity, and (3) overexpression of TgImpα rescues parasites. Although it is tempting to speculate that the weak auto-inhibition of TgImpα might allow binding of the SV40-NLS at lower concentrations, our results do not support this. Mutants of TgImpα that show stronger auto-inhibition or a lack of auto-inhibition also rescued parasites from NLS toxicity.

However, the strength of auto-inhibition of TgImpα is increased or decreased by merely two-fold in the mutants tested in this study (Bhambid et al., 2023). If TgImpα could be mutated to generate variants exhibiting much higher auto-inhibition, the SV40-NLS might show lower parasite toxicity. Literature supports this indirectly: expression of SV40-NLS-GFP constructs in human and *S. cerevisiae* cells, where the Impα proteins show strong auto-inhibition, results in no observable toxicity (Cardarelli et al., 2009; Lobl et al., 1990). Instead, for SV40-NLS to show cell toxicity in *S. cerevisiae*, the strength of SV40-NLS binding to importin α had to be increased to the pM range, generating the Bimax peptides (Kosugi et al., 2008).

The relatively weak auto-inhibition of *T. gondii* importin α may provide a unique advantage as a potential therapeutic strategy. This weak auto-inhibition may allow binding to the wild-type SV40-NLS at lower concentrations, particularly when exposed at the N-terminus of 2xGFP. In contrast, the importin α isoforms of the human host exhibit stronger auto-inhibition, and tight-binding mutants of the SV40-NLS (Bimax peptides) may be required to destabilize the nuclear transport pathway (Kosugi et al., 2008). This is perhaps unexpected since the SV40-NLS is found in a polyomavirus that evolved to hijack the host machinery without large perturbations.

### Possibility of acquiring resistance to the SV40-NLS

Rescuing the lethal phenotype through overexpression of TgImpα proteins confirms that the observed cytotoxicity is mediated via the nuclear import pathway. However, this result suggests parasites may develop resistance by upregulating importin α expression to counteract the toxic effects of the NLS peptide. Our findings show parasite viability can only be restored when importin α is expressed at a sufficiently high level. When stable lines were generated expressing a second copy of TgImpα under the control of the tubulin promoter (ptub1), the expression of this gene was lower (by many orders of magnitude) than that seen for the same expression cassette 22 hours after transient transfection. It appears that *T. gondii* tachyzoites cannot sustain high levels of TgImpα in the long term and tune down expression of the protein, even when driven by a heterologous promoter. Interestingly, rescue of SV40-NLS toxicity was observed only during transient transfections and not in the stable lines.

While these data suggest that resistance to SV40-NLS peptides may not be easily acquired, it would be essential to carry out standard assays using sub-lethal concentrations of the therapeutic molecule (in this case, a cell-permeable NLS peptide) and assess whether parasites can grow after acquiring mutations (Reynolds et al., 2001).

### Delivery of the SV40-NLS into tachyzoites

As the SV40-NLS causes toxicity in tachyzoites, the next step in drug development would involve delivering the peptide to the parasites. For NLS-based peptide inhibitors to be effective, they must efficiently penetrate multiple cellular barriers, including the host cell membrane, the parasitophorous vacuole membrane, and the parasite plasma membrane. Our experiments with one such peptide, the HIV Tat-NLS, did not result in cytotoxicity of tachyzoites; thus, we did not pursue it further. Other peptides, such as oligoarginine, have been reported to transport compounds across mammalian cell membranes, *T. gondii* tachyzoites and bradyzoites membranes, and even across the blood-brain barrier (Futaki, 2005; Futaki et al., 2001; Hao et al., 2022; Jiang et al., 2016; Samuel et al., 2003). Future studies will focus on engineering fusion peptides that combine oligoarginine motifs with the SV40-NLS and evaluating their ability to kill tachyzoites when delivered exogenously.

## CONCLUSION

Peptide inhibitors offer a significant advantage over small molecules by covering larger interaction surfaces and exhibiting stronger binding affinities (Abbasali et al., 2023; Fosgerau & Hoffmann, 2015; Kosugi et al., 2008; Lawrence et al., 2018; Wiedmann et al., 2017). This study is the first to explore the potential of designing NLS peptides to target *Toxoplasma gondii* nuclear import pathways using the SV40-NLS scaffold. This NLS shows remarkable specificity in targeting *T. gondii* tachyzoites while leaving the human host cells unaffected. We envisage future studies focusing on increasing specificity with mutation and delivery strategies and expanding the scope of the therapeutic approach to *Plasmodium* parasites.

## ACKNOWLEDGEMENTS

We thank Vishakha Dey for generating the TgImpα-HA constructs. We acknowledge Sofia Anjum for optimizing the CRISPR-Cas9 strategy and for designing SAG1 and UPRT locus primers utilized in this study. We thank the Indian Institute of Technology Bombay (IIT Bombay) for the central facilities used in this study: the Confocal Laser Scanning Microscope and the Spinning Disc Confocal Microscope. We thank Geetanjali Mishra and Disha Mukherjee for their critical reading of the manuscript.

## FUNDING

The authors acknowledge the International Centre for Genetic Engineering and Biotechnology (ICGEB) for funding to SP (Grant No. CRP/22/005). In particular, we appreciate the efficient and prompt administrative support provided by the ICGEB secretariat. The PhD fellowship from the Human Resource Development Group-Council of Scientific and Industrial Research (HRDG-CSIR) (MB) and the post-doctoral fellowship from IIT Bombay (JC) are acknowledged.

## AUTHOR CONTRIBUTIONS

MB and SP conceived and designed the experiments. MB performed the experiments for all figures, and JC performed the experiments for Figure 1. MB, JC, and SP analyzed the images. MB prepared the figures. MB and SP prepared the manuscript, and JC gave feedback on the drafts. All authors reviewed the final manuscript.

## TRANSPARENCY DECLARATION

The authors declare they have no competing interests.

## SUPPLEMENTARY FIGURES

**Supplementary Figure 1.**
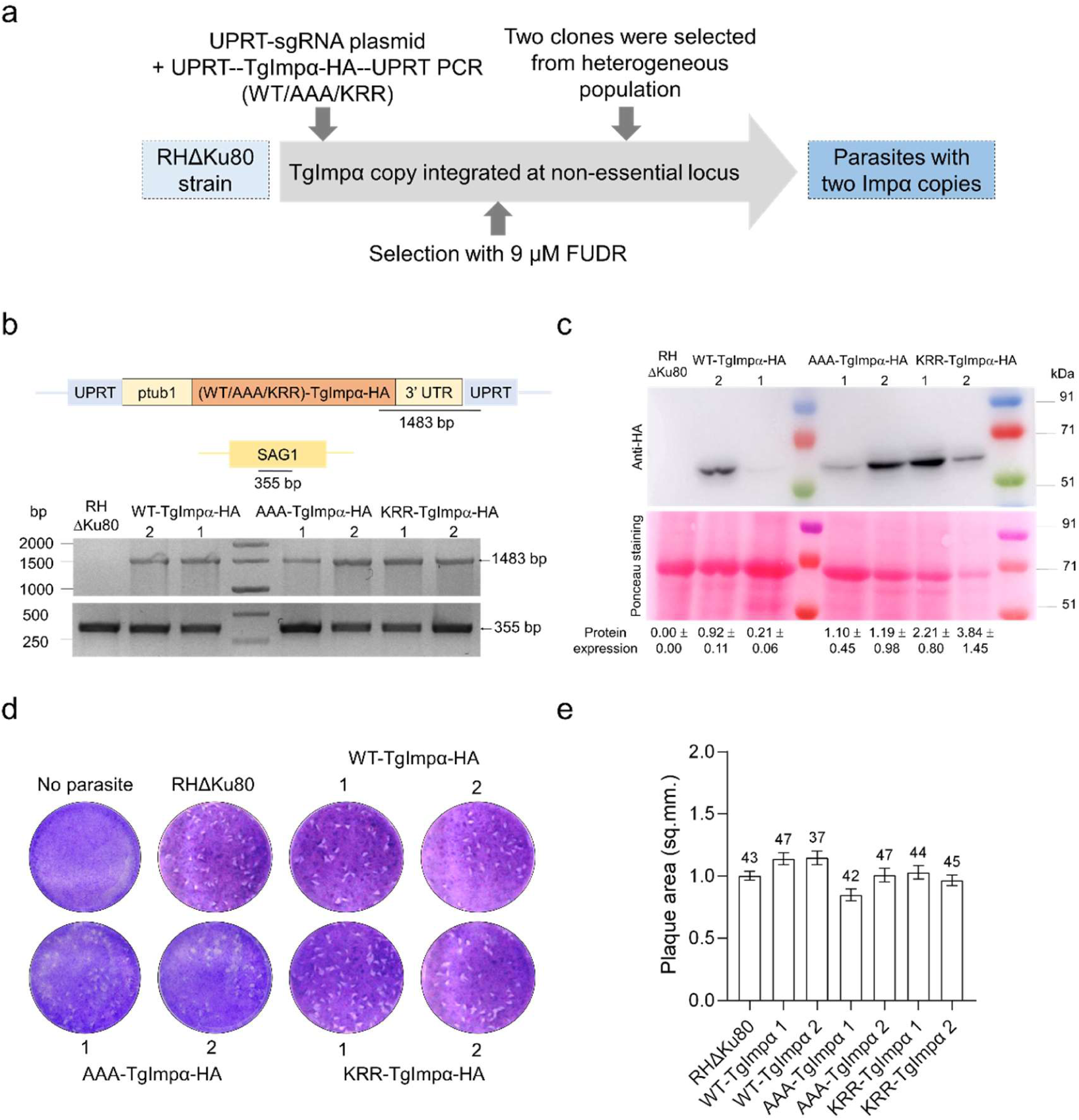
CRISPR/Cas9-mediated integration of TgImpα-HA in the *T. gondii* genome. **(a)** Schematic illustration showing the strategy to incorporate the cassette of TgImpα-HA at the UPRT locus and the selection of transgenic line; **(b)** Schematic of the UPRT and Sag1 locus showing the expected PCR amplicons for screening transfectants. Diagnostic PCR of the UPRT locus yielded a 1.4 kb band demonstrating positive integration (N = 2); **(c)** Expression of wild-type and mutants of TgImpα-HA (60 kDa) determined with Western blot. Ponceau S stained blot was used as a loading control to normalize the protein expression level that is mentioned as mean ± SD (N = 3) in two single plaque clones; **(d)** Plaque assay of the parental line and the stable lines expressing the additional copy of importin α; **(e)** Plaque area was measured, and no significant difference was observed.

**Supplementary Figure 2.**
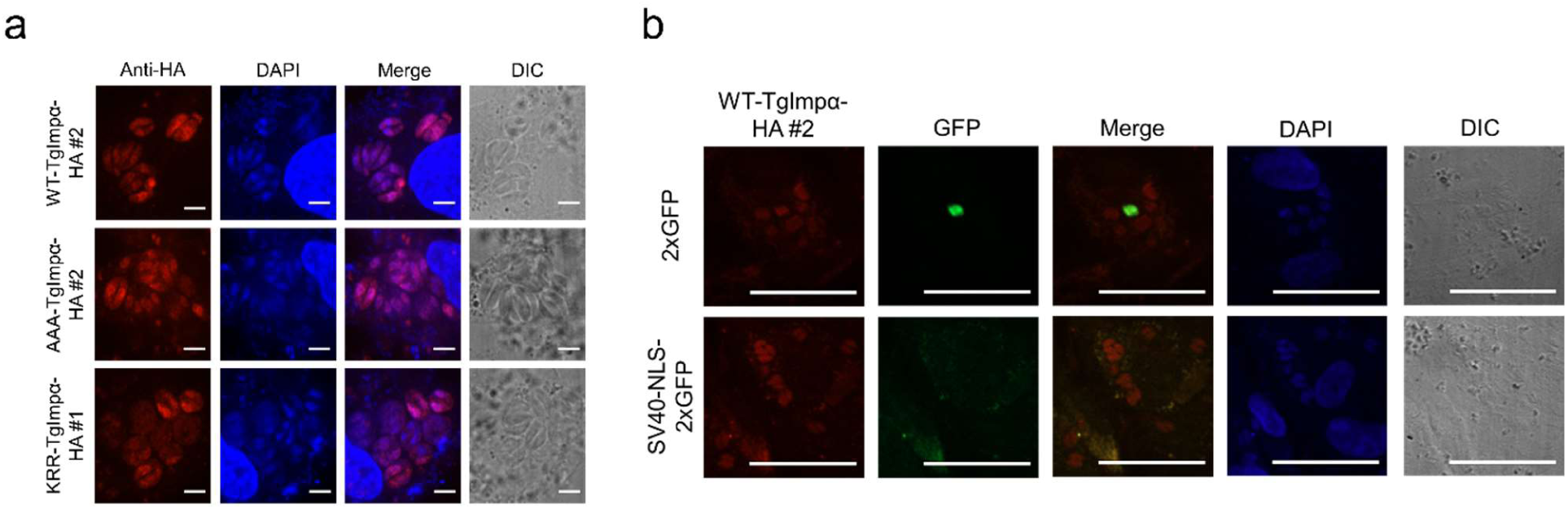
Parasites not rescued from SV40-NLS toxicity in TgImpα stable lines. **(a)** Protein expression was confirmed with immunofluorescence of clonal lines with higher protein levels. Images were captured in a Zeiss LSM 780 confocal microscope with 100X objective (scale bar 5 μm); **(b)** Stable lines of wild-type and variants of TgImpα-HA (red) were transiently transfected with 2xGFP and SV40-NLS-2xGFP (green). Images were captured in a Zeiss Observer Z1 Spinning Disc microscope with a 40X objective (scale bar 50 μm).

## Notes

### Competing Interest Statement

The authors have declared no competing interest.

